# Navigational strategies underlying temporal phototaxis in *Drosophila* larvae

**DOI:** 10.1101/2020.01.06.896142

**Authors:** Maxwell L. Zhu, Kristian J. Herrera, Katrin Vogt, Armin Bahl

**Affiliations:** Department of Physics, Harvard University, Cambridge, MA, USA; Department of Molecular and Cellular Biology, Harvard University, Cambridge, MA, USA; Department of Biology, University of Konstanz, Konstanz, Germany; Centre for the Advanced Study of Collective Behaviour, University of Konstanz, Konstanz, Germany

**Keywords:** *Drosophila* larvae, animal behavior, posture tracking, navigation, phototaxis, modeling

## Abstract

Navigating across light gradients is essential for survival for many animals. However, we still have a poor understanding of the algorithms that underlie such behaviors. Here we develop a novel phototaxis assay for *Drosophila* larvae in which light intensity is always spatially uniform but updates depending on the location of the animal in the arena. Even though larvae can only rely on temporal cues in this closed-loop setup, we find that they are capable of finding preferred areas of low light intensity. Further detailed analysis of their behavior reveals that larvae turn more frequently and that heading angle changes increase when they experience brightness increments over extended periods of time. We suggest that temporal integration of brightness change during runs is an important – and so far largely unexplored – element of phototaxis.

**Summary statement:** Using a novel closed-loop behavioral assay, we show that *Drosophila* larvae can navigate light gradients exclusively using temporal cues. Analyzing and modeling their behavior in detail, we propose that larvae achieve this by integrating brightness change during runs.

## Introduction

Many animals have evolved behaviors to find favorable locations in complex natural environments. Such behaviors include chemotaxis to approach or avoid chemical stimuli; thermotaxis to find cooler or warmer regions; and phototaxis to approach or avoid light (Gepner et al., 2015; Gomez-Marin and Louis, 2014; Gomez-Marin et al., 2011; Kane et al., 2013; Klein et al., 2015; Luo et al., 2010).

*Drosophila* larvae are negatively phototactic, preferring darker regions (Sawin et al., 1994). To navigate, larvae alternate between runs and turns. During runs, larvae move relatively straight. During turns, they slow down and perform head-casts (Lahiri et al., 2011) to sample their environment for navigational decisions (Gomez-Marin and Louis, 2012; Humberg and Sprecher, 2018; Humberg et al., 2018; Kane et al., 2013). However, it is unclear whether such local spatial sampling is necessary to perform phototaxis. Zebrafish larvae, for example, can perform phototaxis even when light intensity is uniform across space but changes over time with the animal’s position (Chen and Engert, 2014). In a purely temporal phototaxis assay, spatial information is absent, so navigation must depend on other cues.

Previous work indicates that as brightness increases, *Drosophila* larvae make shorter runs and bigger turns (Humberg et al., 2018; Kane et al., 2013). This is reminiscent of chemotactic strategies, where decreasing concentrations of a favorable odorant increase the likelihood of turning (Gomez-Marin et al., 2011). While it has been shown that temporal sampling of olfactory cues is sufficient to guide chemotaxis (Schulze et al., 2015), it remains unclear whether larvae can use a purely temporal strategy for visual navigation.

Using a virtual landscape in which brightness is always spatially uniform but depends on the location of the animal in the arena, we confirm that larvae can perform phototaxis by modulating run-length and heading angle. Our data indicate that larvae achieve this by integrating brightness change during runs (**Video S1**).

## Materials and methods

### Experimental setup

All experiments were performed using wild-type 2^nd^-instar *Drosophila melanogaster* larvae collected 3-4 days after egg-laying. This age was chosen to ensure consistent phototactic behavior because older larvae might change their light preference (Sawin-McCormack et al., 1995). Larvae were raised on agarose plates with grape juice and yeast paste, with a 12h/12h light-dark cycle at 22°C and 60% humidity. Before experiments, larvae were washed in droplets of deionized water. All experiments were carried out between 2 pm and 7 pm to avoid potential circadian effects (Mazzoni et al., 2005). Each experiment lasted for 60 min. For all stimuli, animals were presented with constant gray during the first 15 min, allowing them to distribute in the arena.

Larvae were placed in the center of a custom-made circular acrylic dish (6 cm radius) filled with a thin layer of freshly made 2% agarose (Fig. 1A). As previously described (Bahl and Engert, 2020), spatially uniform whole-field illumination was presented via a projector (60 Hz, AAXA P300 Pico Projector) from below. Brightness was set by the computer and ranged from values 0 to 255. Respective light intensity was measured using an Extech Instruments Light Meter LT300 and ranged from 41 Lux to 2870 Lux (**Fig. S1A**). We did not attempt to linearize this curve as it is unclear how the larval visual system processes contrast. Therefore, for all brightness-dependent behavioral analyses, the original pixel brightness value, as set by the program, was used.

**Figure 1.**
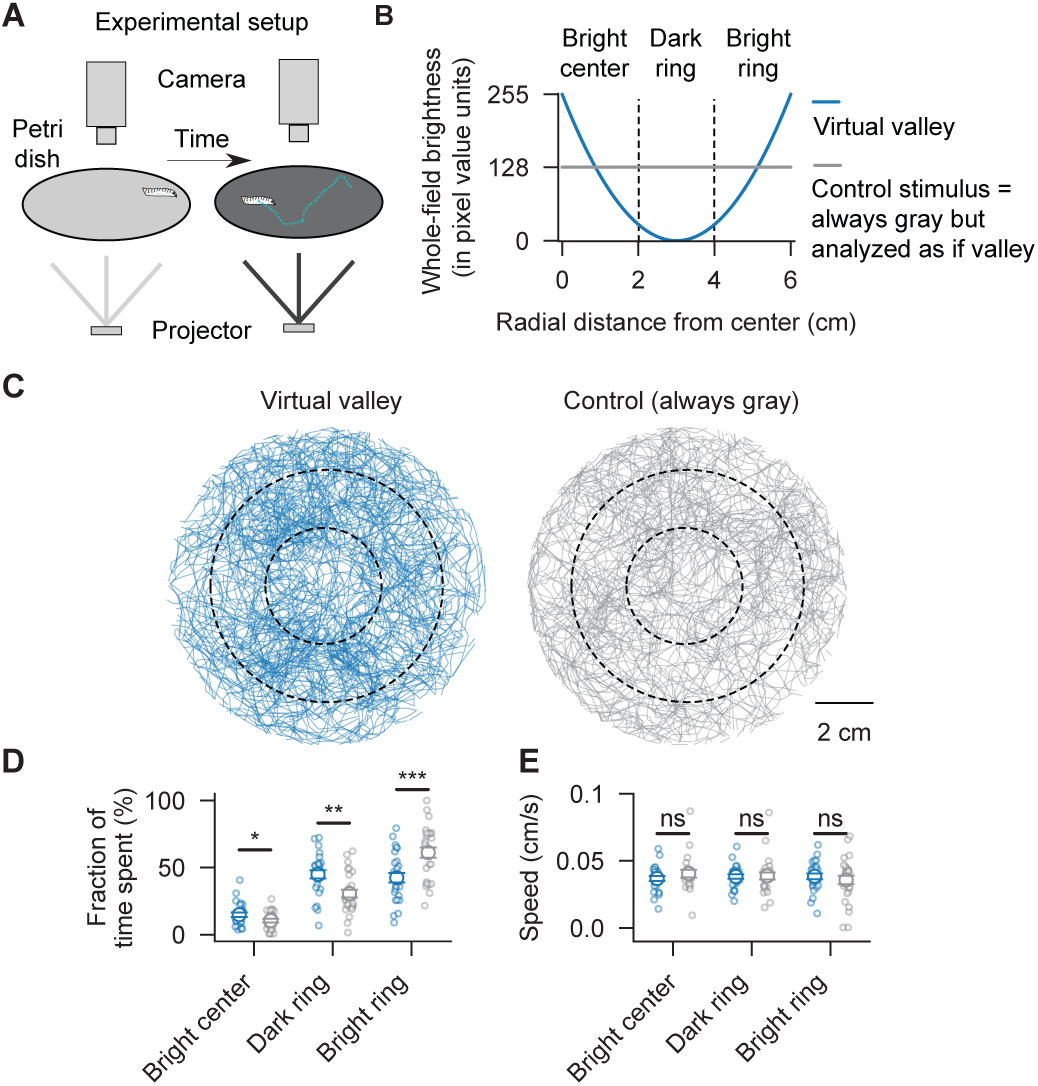
Drosophila larvae can perform temporal phototaxis. (**A**) Setup for tracking freely-crawling *Drosophila* larvae. (**B**) Whole-field pixel brightness versus larval position for the “Valley” and “Control” stimulus. (**C**) Raw trajectories. Dashed circles delineate the “Bright” center, the “Dark” ring, and the “Bright” ring. (**D**) Fraction of time spent in regions (left to right: p = 0.045, p = 0.001, p < 0.001; two-sided t-tests). (**E**) Crawling speed in regions (left to right: p = 0.304, p = 0.891, p = 0.479; two-sided t-tests). Error bars represent mean ± SEM. Blue solid lines and dots indicate “Valley” stimulus larvae; gray solid lines and dots indicate “Constant” stimulus larvae. N = 27 larvae for both groups. Open small circles represent individual animals.

Three virtual light intensity landscapes were tested: a “Valley” stimulus, a “Ramp” stimulus, and a “Constant” stimulus. For the “Valley” and “Ramp” stimuli, the spatially uniform light brightness (*λ*) was updated in closed-loop according to *λ*= 255 · (*r* − 3)^2^/ 9 (Figs. 1B and **S2A**) and 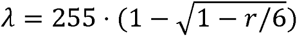 (**Fig. S3A**), respectively, where *r* is the larva’s radial distance to the center of the arena. Both profiles ensure that brightness levels near the wall are high, decreasing the edge preference of larvae and reducing boundary effects. For the “Constant” stimulus, brightness values remained gray (*λ* = 128) regardless of the larva’s position.

For online tracking, the scene was illuminated using infrared LED panels (940 nm panel, 15-IL05, Cop Security). A high-speed camera (90 Hz, USB3 Grasshopper3-NIR, FLIR Systems) with an infrared filter (R72, Hoya) was used to track the larva’s centroid position in real-time. Eight independent arenas were operated in parallel, making the system medium to high-throughput and relatively cost-effective. The position of the animal was determined by spatially filtering the background-subtracted image and then searching for the largest contour. The procedure provides a reliable estimate of the animal’s centroid position but cannot determine the precise location of the head or the tail. Using the centroid as a closed-loop position signal significantly simplifies the experimental procedure and is justified as larvae are small in size relative to the slowly changing and always spatially uniform virtual brightness landscapes. The spatial precision of our tracking was in the order of ±0.01 cm per ∼10 ms, resulting in a nearly noise-free presentation of the stimulus profiles (**Fig. S1B**). In addition to the online-tracking, a video of the animal was stored for offline posture analysis (**Video S2**).

In our system, the closed-loop latency between the detection of the animal’s position and the update of the visual stimulus is 100 ms. This value was determined using the following protocol: Infrared filters were removed from the cameras, allowing for direct measurements of the brightness from the projector. Arena brightness starts at a high level but is set to a dark state after a few seconds. When the camera detects such an event, the computer sets the brightness back at a high level. The length of the resulting dark period is the closed-loop delay. Using this strategy, the resulting value contains the sum of all delays of the system (camera image acquisition, image buffering, data transport to the USB 3.0 hub, PCI-express to CPU transport, CPU image analysis, command to the graphics card, graphics buffering, and buffering and image display on the projector). While it is hard to use GPU-based systems to reach closed-loop delays below 100 ms (Stowers et al., 2017), simpler systems with direct LED control allow for delays as short as 30 ms (Tadres and Louis, 2020).

### Control experiments

Notably, animals navigating the “Constant” stimulus were always analyzed as if they navigated the respective experimental stimulus (“Valley” or “Ramp”), using the same binning, naming conventions, and analysis methods. For example, control animals that spend time in the “Dark” ring (gray open circles in Fig. 1D) actually perceive constant gray during the entire experiment. This analysis was chosen to control for the spatial arrangement of our stimulus and boundary effects. The best example where this strategy is important can be seen for the turn-triggered brightness change (Fig. 2G): Even though control animals always perceive gray, the turn-triggered brightness dynamics indicate a complex dependency on the spatial arrangement of the arena. Only by using this control analysis is it possible to appreciate the dynamics in the experimental group.

**Figure 2.**
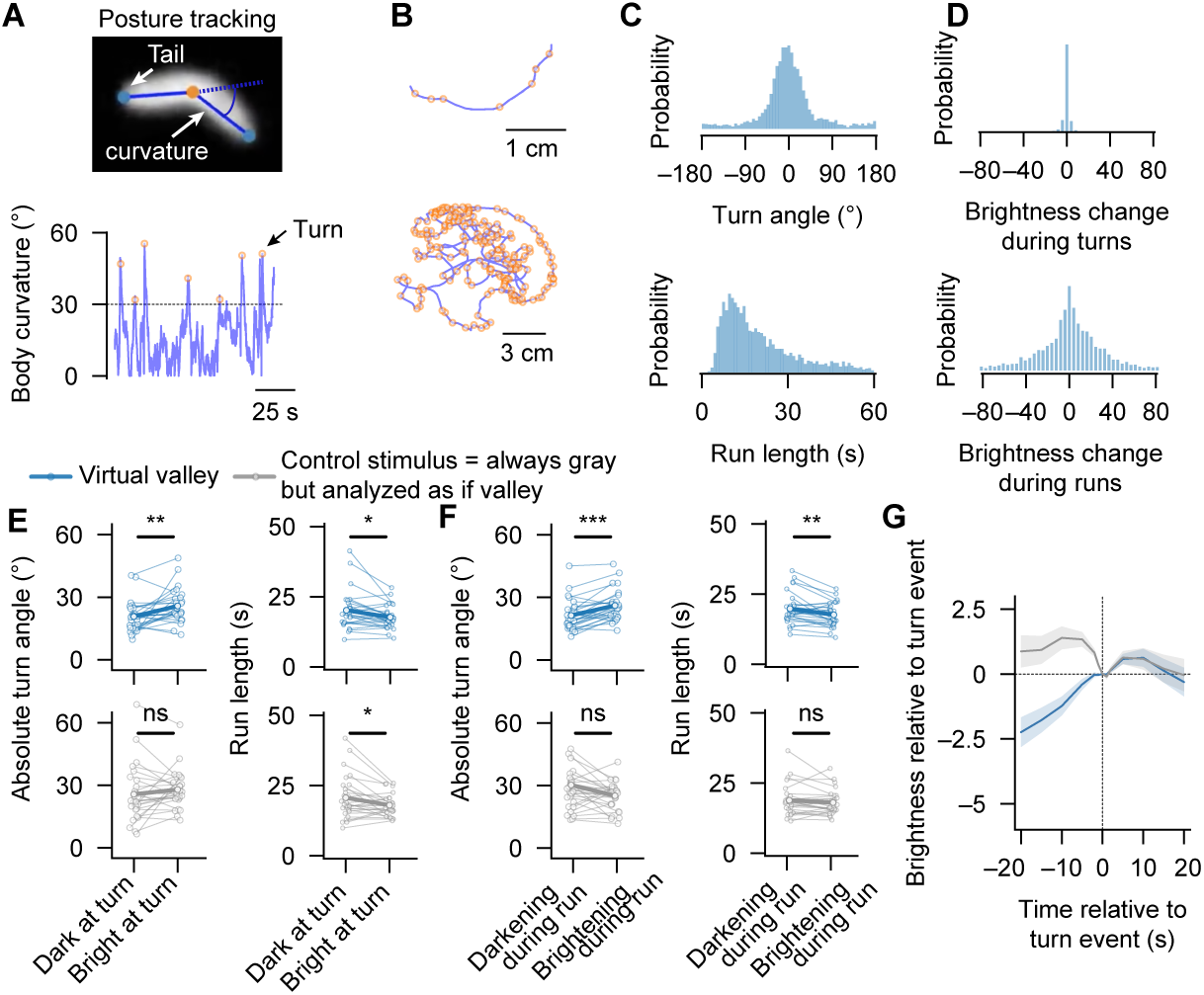
Brightness and brightness history modulate navigational decisions. (**A**) Posture tracking for estimating larval body curvature (angle between solid and dashed blue lines). Turns (orange circles) are curvature peaks above a threshold (30°). (**B**) Example trajectory with detected turns for an inset view (top) and the entire arena (bottom). (**C,D**) Probability density distributions for turn angles and run-length (**C**) and respective brightness changes (**D**). (**E,F**) Turn angle and run-length as a function of light intensity (dark: < 29; bright: otherwise; see brightness profile, Fig. 1B) and as a function of brightness change since the previous turn (left to right: p = 0.004, p = 0.010, p < 0.001, p = 0.006 for the “Valley” stimulus and p = 0.289, p = 0.018, p = 0.066, p = 0.221 for the “Constant” stimulus; paired t-tests). (**G**) Turn event-triggered brightness for the “Valley” and the “Constant” stimulus (mean ± SEM over all turns from all larvae, n = 3153 and n = 2981 turns, respectively). N = 27 larvae for both groups. Open small circles and thin solid lines in (**E,F**) represent median turn angle and run-length for individual larvae.

### Data analysis and statistics

All data analysis was performed using custom-written Python code on the 45 min period after acclimatization. To avoid tracking problems and minimize boundary effects, data were excluded where larvae were within 0.1 cm distance to the edge.

The circular arena was binned in three concentric regions depending on the radius *r:r =* 0 − 2 *cm, r =* 2 − 4 *cm,* and *r =* 4 − 6 *cm.* These regions were named the “Bright” center, the “Dark” ring, and the “Bright” ring for the “Valley” stimulus (Fig. 1B) and the “Dark” center, the “Gray” ring, and the “Bright” ring for the “Ramp” stimulus (**Fig. S3A**). Animal speed was computed by interpolating the trajectory to 1 s bins and then by taking the average distance of consecutive points (Fig. 1E).

For the turn event-based offline analysis (Fig. 2), a pose estimation toolbox, DeepPoseKit (Graving et al., 2019), was used. To this end, 100 frames were manually annotated (head, centroid, and tail) to train the neural network, which was then used to predict animal posture across all frames from all animals. Body curvature was defined as the angle between the tail-to-centroid vector and the centroid-to-head vector (Fig. 2A). The pose estimation algorithm occasionally had difficulties distinguishing between the head and the tail. This problem was, however, not relevant for the curvature measurement as the angle between these two body parts does not change when they are flipped. In a few frames, the algorithm placed the head and the tail at the same location, leading to the transient detection of large body curvatures. These events were discarded by low-pass filtering traces with a Butterworth filter (cutoff frequency: 3 Hz). Turn events were defined as a local curvature peak above 30° and needed to be separated from the previous event by at least 2 s in time and 0.2 cm in space. The value for the curvature threshold was chosen such that the identified curvature peaks clearly stood out from the curvature fluctuations in between events (Fig. 2A).

Turn angles were defined as the angle between the location in the arena 2 s before a turn event and 2 s after. Run-length was defined as the time between consecutive turn events. Each turn event was labeled as “Dark” or “Bright”, based on the brightness equations and binning described above (Dark: pixel brightness less than 29, Bright: otherwise), and as “Darkening” or “Brightening” based on the sign in brightness change since the last turn event (Fig. 2E,F). As turn events are short and spatially confined, by stimulus design, the whole-field brightness change during such events is nearly zero (Fig. 2D). Notably, our curvature-based turn event identification procedure does not allow for precise labeling of the beginning and the end of the event. Therefore, the brightness change during turns was defined as the brightness difference 0.5 s before and 0.5 s after the event. This time range often includes brief periods of runs, explaining the small residual width of the reported brightness distribution (Figs. 2D and 3E). The brightness change during runs was defined as the difference in brightness between two consecutive turn events (Figs. 2D and 3E).

**Figure 3.**
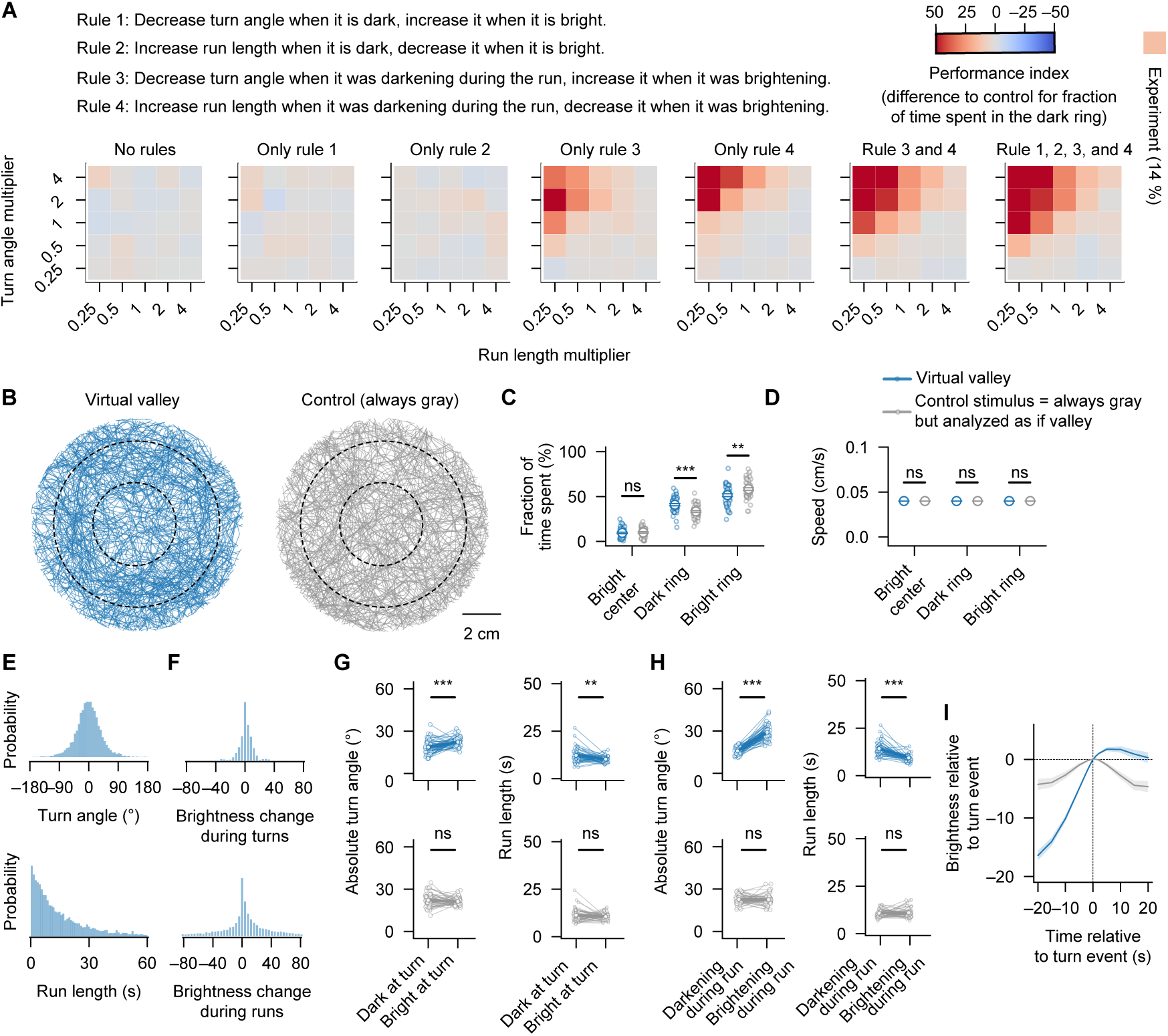
Simulated larvae perform temporal phototaxis. (**A**) Characterization of combinations of four potential navigational rules, with a grid search for the parameters run-length and turn angle, quantified by a phototaxis performance index. (**B-I**) Simulations using only Rules 3 and 4, with turn angle and run-length multiplier set to one. (**B-D**) Raw trajectories, fraction of time spent in regions, and crawling speed (as in Fig. 1C-E). Left to right: (**C**) p = 0.181, p < 0.001, p = 0.015; two-sided t-tests; (**D**) p = 0.531, p = 0.651, p = 0.665; two-sided t-tests. (**E-I**) Analysis of turns and runs (as in Fig. 2C-G). (**G,H**) Left to right: p < 0.001, p = 0.001, p < 0.001, p < 0.001 for the “Valley” stimulus; p = 0.283, p = 0.165, p = 0.796, p = 0.656 for the “Constant” stimulus; paired t-tests. Open circles and thin solid lines in (**C-I**) represent individual model larvae. N = 50 simulation runs for both groups using different random seeds. N = 5331 and n = 5334 events in (**I**).

Two-sample t-tests were used for pairwise comparisons between the experimental and control data. Paired-sample t-tests were used for pairwise comparisons within groups. Statistics for the linear regression fits (**Figs. S4A,B** and **S5A,B**) were based on a bootstrapping approach by repeating the analysis 1000 times for shuffled data and then comparing the distribution of R^2^ values to the one from the original dataset.

Larvae were discarded if they spent more than 99% of the experimental time in a single region or if their speed was zero. All data analysis was done automatically in the same way for the experimental and control groups.

### Modeling

Simulations (Figs. 3, **S5, and S6**) were custom written in Python 3.7, using the high-performance Python compiler numba. Simulations were performed using Euler’s Method with a timestep of dt = 0.01 s. Model larvae were initialized with a random position and orientation. At each time step, larvae stochastically chose one of two possible actions: They could either move forward, with a speed of 0.04 cm/s (parameter was taken directly from the experiment, Fig. 1E), or turn. The baseline probability for turning was p = 0.00066. This value was directly computed from the experiment to match the measured average run-length of T = 15 s (Fig. 2E,F), following p = dt / T. When making turns, turn angles were drawn from a Gaussian distribution with a baseline standard deviation of 32°, matching the experimental value (Fig. 2C,E,F). When model larvae reached the edge, a new random direction vector was chosen, preventing them from leaving the arena.

In correspondence with our experimental findings (Fig. 2E,F), the model was equipped with four additional navigational rules (Fig. 3A).

“Rule 1”: When the environment is “Dark” (brightness smaller than 29), turn angles decrease. When it is “Bright” (brightness larger than 29), turn angles increase.

“Rule 2”: When the environment is “Dark” (brightness smaller than 29), run-lengths increase. When it is “Bright” (brightness larger than 29), run-lengths decrease.

“Rule 3”: When the environment is “Darkening” (change since previous turn smaller than zero), turn angles decrease. When it is “Brightening’’ (change since previous turn larger than zero), turn angles increase.

“Rule 4”: When the environment is “Darkening” (change since previous turn smaller than zero), run-lengths increase. When it is “Brightening’’ (change since previous turn larger than zero), run-lengths decrease.

Changes in turn angle were accomplished by adjusting the standard deviation of the Gaussian distribution by ±30%, the effect size observed in our experiments (Fig. 2E,F). We modulated run-length (T) by scaling them by ±30%, thereby modulating the probability of turning (p = dt / T). When combinations of those rules were tested (Fig. 3A), effects were concatenated.

A performance index (PI) (Fig. 3A) was used to characterize how well animals or models performed temporal phototaxis. The metric was based on the difference between the experimental and control group for the fraction of time spent in the “Dark” ring. To compute this value, bootstrapping was used to average 1000 samples of randomly chosen differences between experimental and control conditions.

For the parameter grid search (Fig. 3A), the absolute turn angle and the run-length were varied systematically. To this end, respective baseline parameter values (taken from the experiment, Fig. 2E,F), were changed by scaling them with two multipliers (run-length multiplier and turn angle multiplier).

Data generated from model larvae were analyzed and displayed using the exact same scripts that were used to analyze experimental data, allowing for easy comparison between model and animal behavior.

## Results

### Fly larvae can navigate a virtual brightness gradient

We first asked whether fly larvae can perform temporal phototaxis, i.e. navigate a virtual light landscape lacking spatial information. We placed individual animals in an agarose-filled arena, allowed them to freely explore, and tracked their position in real-time (Fig. 1A). We presented spatially uniform light from below, with brightness levels following a quadratic dependence of the larva’s distance from the center (“Valley” stimulus, Fig. 1B) or constant gray as a control (“Constant” stimulus). For both groups, we analyzed how animals distribute across three concentric regions: the “Bright” center, the “Dark” ring, and the “Bright” ring. Notably, throughout this study control animals were always analyzed as if they navigated the experimental stimulus even though they in fact perceived constant gray. This analysis is important to control for the spatial arrangement of our stimulus and boundary effects.

Larvae that navigated the “Valley” stimulus spent a significantly higher fraction of time in the “Dark” ring than those that navigated the “Constant” stimulus (Figs. 1C,D and **S2B**). This behavior was most pronounced between minutes 10 and 40 of the experiment (**Fig. S2C**). To verify that this behavior was not an artifact of our specific stimulus design, we also tested a gradient where brightness monotonically “ramps” with radial distance (**Fig. S3A**) and observed that larvae also here navigated to dark regions (**Fig. S3B,C**).

Because larvae lacked spatial brightness cues in our setup, it was unclear which behavioral algorithms they employ. One basic, yet potentially sufficient, algorithm would be to reduce movement in darker regions. However, speed was independent of brightness (Figs. 1E and **S3D**), suggesting that larvae employ more complex navigational strategies.

We conclude that *Drosophila* larvae are capable of performing phototaxis in the absence of spatial information and that this behavior cannot be explained by a simple brightness-dependent modulation of crawling speed.

### Larval temporal phototaxis depends on brightness change over time

In spatially differentiated light landscapes, fly larvae make navigational decisions by sampling brightness differences during head-casts. In our setup, by design, larvae experience no brightness fluctuations during head-casts. Hence, they have to use whole-field brightness or brightness history information to modulate the magnitude and/or frequency of turns. To explore this possibility, we segmented trajectories into runs and turns. We applied a deep learning-based package, DeepPoseKit (Graving et al., 2019) to extract the larvae’s head, centroid, and tail positions from the experimental video (Fig. 2A and **Video S2**). From there, we calculated the animal’s body curvature to identify head-casting events and to quantify turn angles and run-lengths (Fig. 2A-C).

As expected, brightness changes during the spatially confined turns were negligible compared to ones measured during runs (Fig. 2D). To quantify the effect of brightness on heading angles and run-lengths, we checked how these parameters varied with the larva’s position. During the “Valley” but not the “Constant” stimulus, turns in the “Dark” region led to smaller heading angle changes than in the “Bright” regions (Fig. 2E). Similarly, runs before a turn in the “Dark” region of the “Valley” stimulus were slightly longer compared to runs ending in the “Bright” region. However, this also partly occurred with the “Constant” stimulus, suggesting that the effect might not arise from a visuomotor transformation.

Next, we explored whether brightness history affects behavior. As run-lengths were highly variable, ranging from ∼3 s to ∼40 s (Fig. 2C), we focused our analysis on the brightness change between consecutive turns. We classified turns by whether larvae experienced a decrease or increase in whole-field brightness during the preceding run. We found that heading angle changes were smaller and that run-lengths were longer when larvae had experienced a brightness decrease compared to an increase (Fig. 2F). We did not observe these effects in control animals.

To further quantify the effects of brightness and brightness change on heading angle change, we performed regression analysis directly on individual events (**Fig. S4**). While turn angles scale with brightness, they do so more strongly with brightness change.

These observations led us to hypothesize that larvae might integrate information about the change in brightness during runs and that this integration period might span several seconds. To obtain an idea about time-scales, we computed a turn event-triggered brightness average (Fig. 2G). We observed that, on average, turns performed in the “Valley” stimulus are preceded by an extended period of >20 seconds of brightening, suggesting that long-term brightness increases drive turns.

In summary, our analysis of turns and runs confirms that, first, brightness levels modulate heading angle change and, second, changes in brightness prior to turns modulate heading angle change as well as run-length.

### A simple algorithmic model can explain larval temporal phototaxis

We next wanted to test whether the identified behavioral features are sufficient to explain larval temporal phototaxis. Based on our experimental findings (Fig. 2), we propose four rules as navigational strategies (Fig. 3A). For rules 1 and 2, the instantaneous brightness modulates the heading angle change and run-length, respectively. By contrast, for rules 3 and 4, the brightness change since the last turn modulates the heading angle changes and run-lengths.

To test these navigational rules, we simulated larvae as particles that could either move straight or make turns. To compare the performances of different models, we calculated a phototaxis index (difference of time spent in the “Dark” ring between experimental and control groups, Fig. 3A). For all permutations of our rules, we explored a set of multipliers for the heading angle change and run-length, with a multiplier of 1 corresponding to the experimental averages (Fig. 2E,F). This allowed us to assess the robustness of our model to parameter choice. As expected, with no active rules, the larval distribution was comparable between the “Valley” and “Constant” stimulus. Activating rules 1 or 2, performance did not improve, suggesting that modulation of behavior based on instantaneous brightness is insufficient to perform temporal phototaxis. Activating rules 3 or 4, phototaxis emerged for small run-lengths and large turn angle multipliers. However, for multipliers set to 1, the resulting phototaxis index was weaker than in experiments (= 14 %). Only when combining rules 3 and 4, phototaxis performance matches the experimental values. Combining all four rules yielded minimal improvements. Therefore, for further analysis, we focused on a combination of rules 3 and 4, with both multipliers set to 1.

Like real larvae (Fig. 1C-E), simulated larvae navigating the “Valley” stimulus spent more time in the “Dark” ring than larvae navigating the “Constant” stimulus (Fig. 3B,C) without modulating speed (Fig. 3D). Furthermore, distributions of turn angle changes, run-lengths, and brightness changes were comparable to experimental data (compare Figs. 2C,D and 3E,F). Residual differences in those distributions are likely due to additional mechanisms used by the animal, such as a refractory period for turn initiation, which we did not incorporate in our model. When we examined the effects of instantaneous brightness and brightness change on turn angle amplitude and run-length (Fig. 3G,H), we observed similar patterns as in the experimental data (Fig. 2E,F). As found in experiments (Fig. 2G), turns are preceded by long stretches of increasing brightness (Fig. 3I), supporting our hypothesis that larvae integrate brightness change over several seconds. Moreover, in the event-based regression analysis we found results to be in agreement with experimental data as well (compare **Figs. S4** and **S5**). Finally, to verify that our model generalizes to other visual stimulus patterns, we simulated larvae exploring the “Ramp” stimulus and observed phototaxis performance comparable to that of real larvae (compare **Figs. S3** and **S6**).

In summary, after implementing our experimentally observed navigational rules in a simple computational model, we propose that the most critical element of larval temporal phototaxis is the ability to integrate brightness change over extended time periods. Modulating turn angle amplitude and run-length based on such measurement is sufficient to perform temporal phototaxis.

## Discussion

Using a closed-loop behavioral assay, we show that *Drosophila* larvae find the darker regions of a virtual brightness gradient that lacks any spatial contrast cues. Temporal phototaxis behavioral algorithms have already been dissected in open-loop configurations, where stimuli are decoupled from an animal’s actions. Following a global brightness increase, larvae are known to modify both their heading angle magnitude and their run-length (Gepner et al., 2015; Kane et al., 2013), which is in agreement with our findings. We were able to demonstrate that these navigational strategies are in fact sufficient for phototactic navigation. Given that brightness fluctuations in our assay are slow and negligibly small during head-casts, we suggest that animals integrate brightness change during runs to make decisions about the strength and timing of turns. Previous work has shown that larvae can navigate olfactory or thermal gradients using only temporal cues (Luo et al., 2010; Schulze et al., 2015). Together with our findings, this should enable future exploration of the shared computational principles and neural pathways across these sensory modalities.

Closed-loop systems are powerful tools to dissect an animal’s sensorimotor transformation. They have been employed in many models including adult *Drosophila* (Bahl et al., 2013), larval zebrafish (Bahl and Engert, 2020; Chen and Engert, 2014), and *C. elegans* (Kocabas et al., 2012; Leifer et al., 2011). Recent work in *Drosophila* larvae used LED-based devices to study closed-loop temporal chemotaxis in virtual optogenetic environments (Tadres and Louis, 2020). Such systems are cheaper and have shorter stimulus refresh times but cannot easily be used to present animals with spatially differentiated landscapes. By utilizing a projector, our setup overcomes this limitation. With the drawback of slightly longer delays and higher component costs, the ability to present any type of visual stimulus adds important flexibility and versatility.

Future studies could use our paradigm to study, for example, specific behavioral differences between animals navigating a true luminance gradient compared to when they navigate the exact same one virtually. Moreover, our system makes it possible to explicitly investigate navigational strategies exclusively using spatial information. This has already been achieved in zebrafish larvae (Chen et al., 2020; Huang et al., 2013) by always locking a sharp contrast edge to the center of the animal’s head. Testing such stimuli in *Drosophila* larvae will, however, require more precise real-time position, orientation, and posture measurements, improvements that can be added to our setup. The result from such experiments could be used to construct a spatial phototaxis model which could then be combined with our proposed temporal phototaxis model.

## Acknowledgments

We thank L. Hernandez-Nunez for discussions and reading through the manuscript. We are grateful to F. Engert and A. Samuel and their lab members for discussions and general support.

## Author contributions

All authors contributed equally to the design of the project. A.B. built the behavioral setup. M.Z. performed experiments. M.Z. and A.B. and analyzed data. M.Z., K.J.H., K.V., and A.B. wrote the manuscript. K.J.H., K.V., and A.B. supervised the work.

## Data availability

The data that support the findings of this study are available from the corresponding author upon request. Source code for data analysis and modeling are available on GitHub (https://github.com/arminbahl/drosophila_phototaxis_paper).

