## supplementary information for "Navigational strategies underlying temporal phototaxis in *Drosophila* larvae"

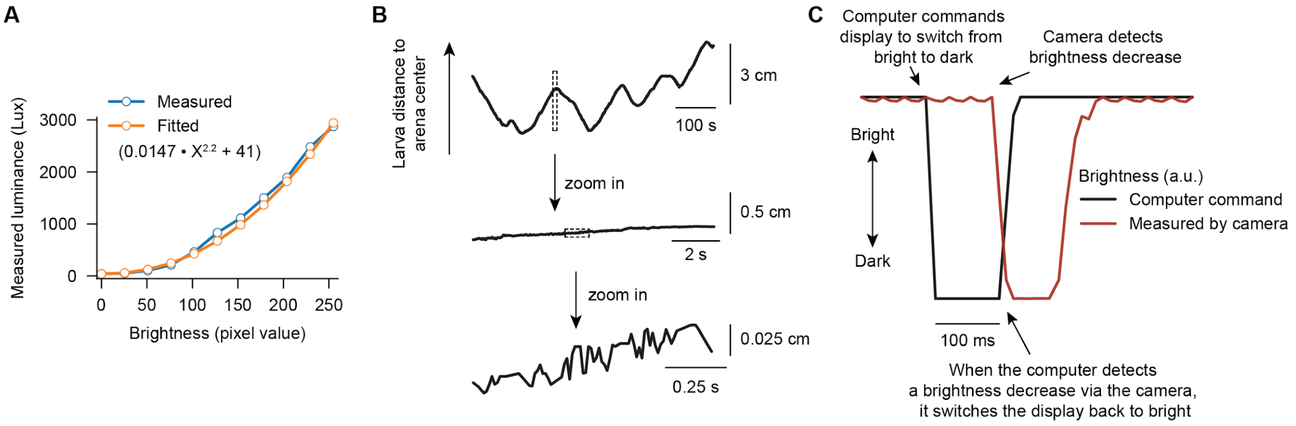


**Figure S1. Performance of the closed-loop behavior setup.** (**A**) Luminance as a function of computer pixel brightness, measured with a professional light meter. Fitting this function with a simple polynomial shows a nearly quadratic relationship. As it is unclear how the larval visual system processes contrast, we did not attempt to linearize this curve. (**B**) Example raw tracking data, displayed in form of the radial distance to the center of the arena. Changes in the slope of this curve indicate events where the larva makes a turn. When zooming into this trace, it becomes visible that tracking noise is very small, in the order of ±0.025 cm (within a period of 0.25 s). Given our position-dependent brightness function (**Fig. 1B**), this would lead to a maximum pixel brightness fluctuation of as small as ±3 over the same short time period. (**C**) Measurement of the closed-loop delay of the system: We remove the infrared filters from the camera and directly measure the luminance coming from the projector. The projector starts with a high level of brightness. After a few seconds, it rapidly switches to a dark state. Whenever the camera detects this event, we automatically set the brightness level back to a bright state again. The time of darkness is the full closed-loop delay of the system.

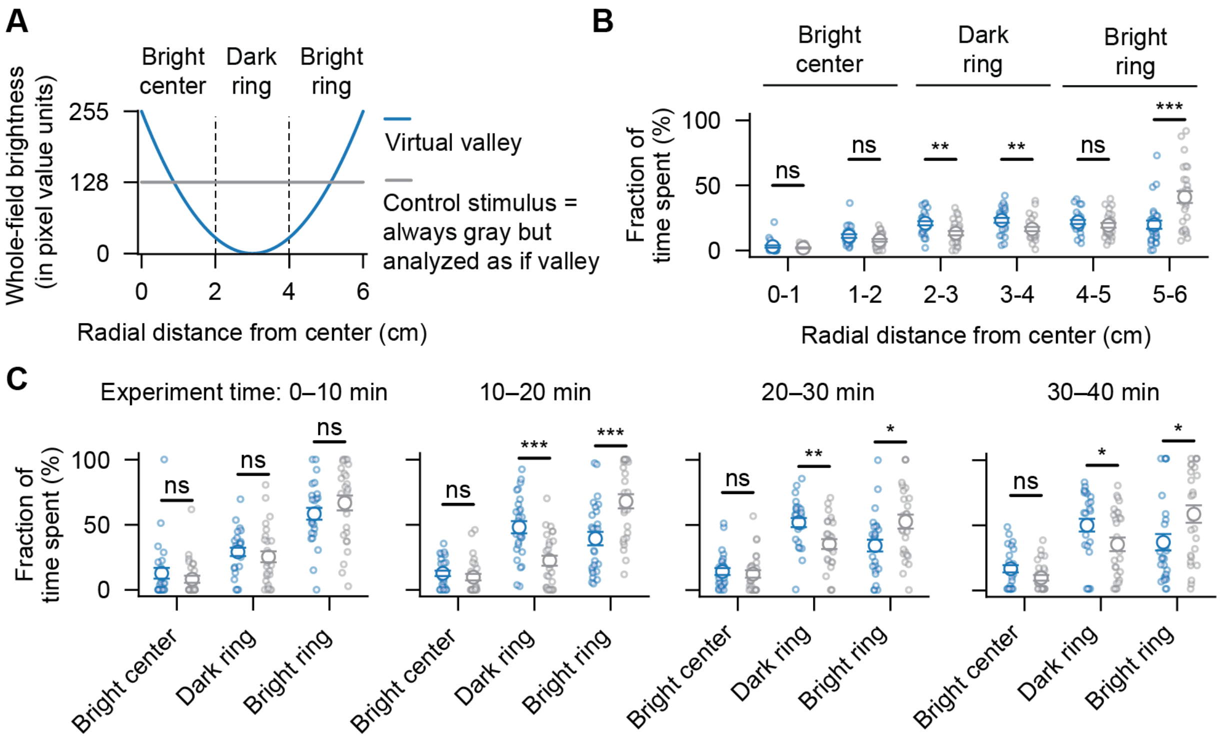


**Figure S2. Detailed analysis of behavioral performance during temporal phototaxis. (A)** Luminance profile used for the "Valley" stimulus (same panel as in **Fig. 1B**). (**B**) Fraction of time larvae spend in the different regions for finer radial bin size (left to right: $p = 0.264$, $p = 0.060$, $p = 0.002$, $p = 0.009$, $p = 0.310$, $p < 0.001$; two-sided t-tests). (**C**) Fraction of time spent in the different regions, analyzed in 10 min time bins (left to right: $p = 0.360$, $p = 0.481$, $p = 0.256$ for first period; $p = 0.053$, $p < 0.001$, $p < 0.001$ for second period; $p = 0.616$, $p = 0.002$, $p = 0.011$ for third period; $p = 0.054$, $p = 0.044$, $p = 0.022$ for third period; two-sided t-tests). Error bars in represent mean ± SEM. Blue dots in indicate "Valley" stimulus larvae. Gray dots indicate "Constant" stimulus larvae. N = 27 larvae for both groups. Open circles represent individual animals.

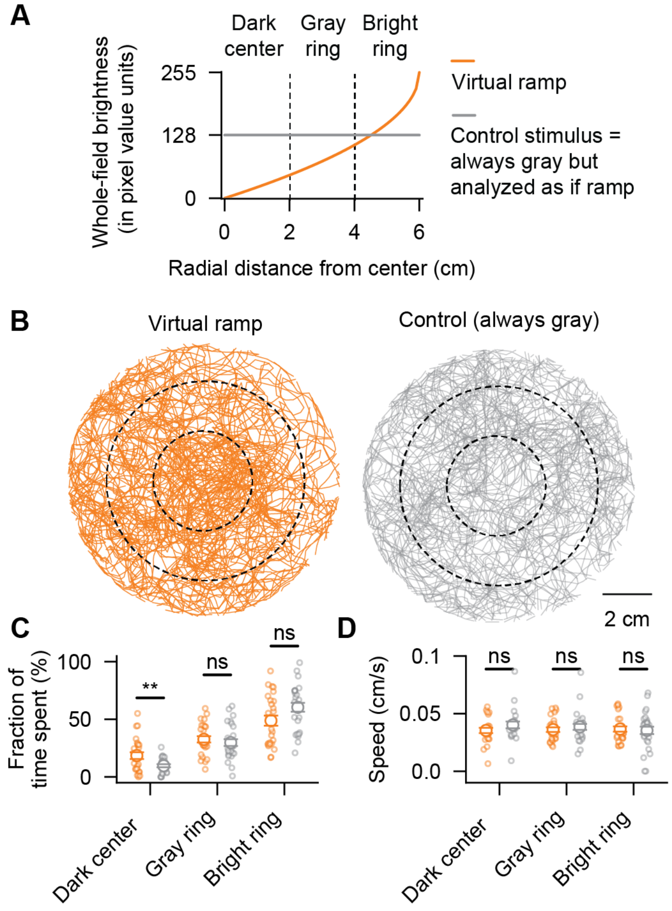


**Figure S3. Temporal phototaxis in the alternative "Ramp" stimulus configuration.** (**A**) Whole-field brightness as a function of larval position for the "Ramp" stimulus and the "Control" stimulus. (**B**) Raw trajectories. Dashed circles delineate the "Dark" center, the "Gray" ring, and the "Bright" ring. (**C**) Fraction of time spent in each region (left to right: p = 0.009, p = 0.484, p = 0.051; two-sided t-tests). (**D**) Crawling speed in each region (left to right: p = 0.200 p = 0.479, p = 0.770; two-sided t-tests). Error bars represent mean ± SEM. Orange dots and lines indicate "Ramp" stimulus larvae; Gray dots and lines indicate "Constant" stimulus larvae. N = 26 and N = 27 larvae for the "Ramp" stimulus and the "Constant" stimulus, respectively. Open circles represent individual animals.

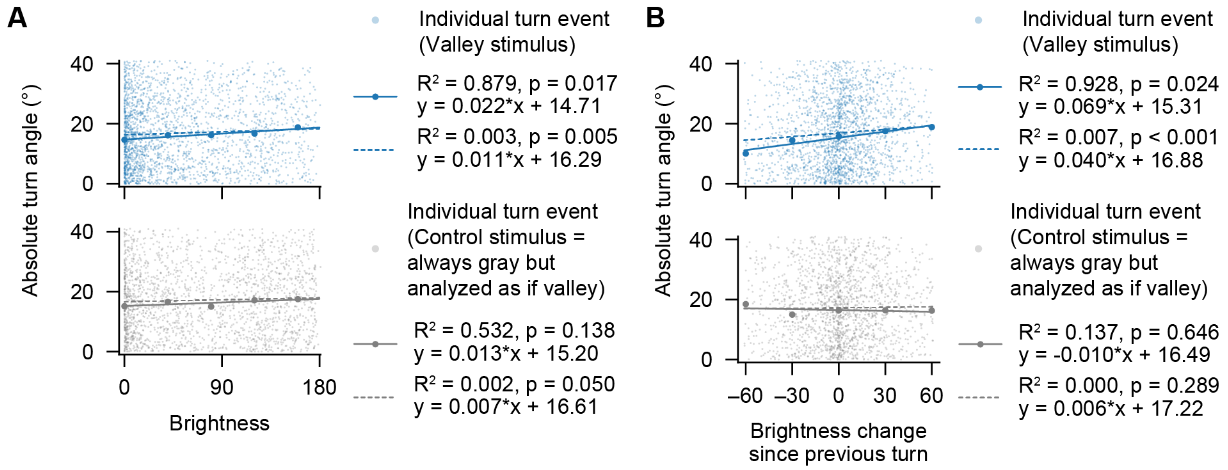


**Figure S4. Regression analysis on individual turn events (Experimental data). (A,B)** Absolute turn angle as a function of instantaneous brightness (**A**) and as a function of brightness change since the previous turn (**B**) for “Valley” stimulus larvae (blue) and “Constant” stimulus larvae (gray). Small dots represent individual turns (n = 3153 and n = 2981 events, respectively, from N = 27 larvae for each group). Bigger filled dots are binned data. Regression analysis was performed on binned data (solid lines) as well as on raw data points (dashed lines). P values are based on a bootstrapping approach by repeating the regression analysis 1000 times for shuffled raw data and then comparing the distribution of $R^2$ values to the one from the original dataset.

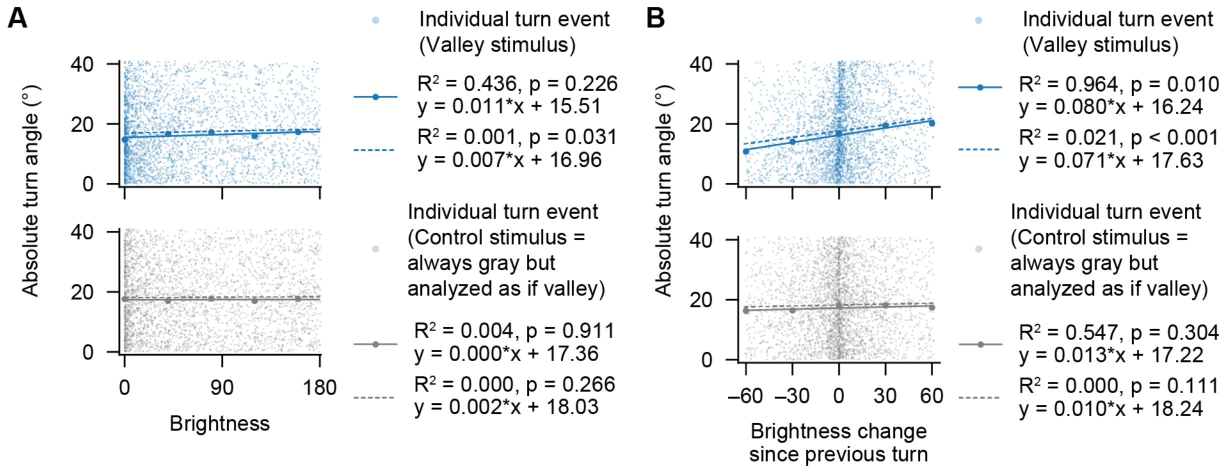


**Figure S5. Regression analysis on individual turn events (Model data). (A,B)** Same analysis and statistics as in **Fig. S4A,B** but for model data.

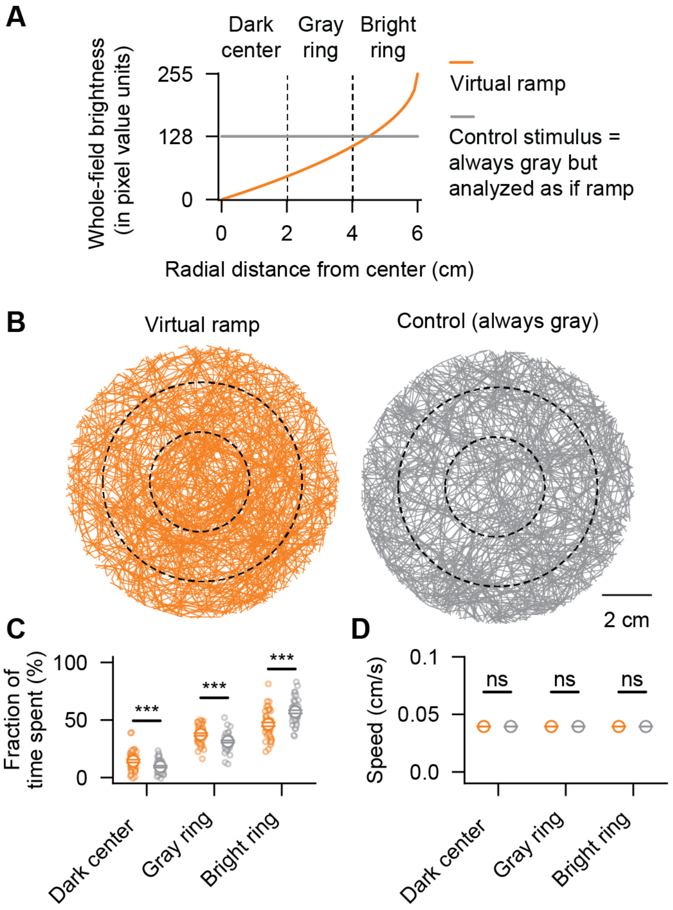


**Figure S6. Simulated larvae navigating the "Ramp" stimulus. (A)** Whole-field brightness as a function of larval position for the "Ramp" stimulus and the "Control" stimulus (same as in **Fig. S3A**). (**B**) Raw trajectories for model larvae navigating the "Ramp" stimulus (orange) and the "Constant" stimulus (gray). Dashed circles delineate the "Dark" center, the "Gray" ring, and the "Bright" ring. (**C,D**) Fraction of time simulated larvae spend in the different regions and respective crawling speeds (as in **Fig. S3C,D**); left to right: $p < 0.001$, $p < 0.001$, $p < 0.001$ in (**C**) and $p = 0.296$, $p = 0.677$, $p = 0.213$ in (**D**). N = 50 simulated larvae in both groups. Model parameters are the same as used in **Fig. 3B–I**. Error bars represent mean ± SEM.
